# THE PHYTOPHARMACOLOGICAL EVALUATION OF THE METHANOL ROOT EXTRACT OF *AZANZA GARCKEANA* (MALVACEAE) ON ISOLATED WISTAR RAT UTERINE SMOOTH MUSCLES

**DOI:** 10.1101/2024.05.10.593648

**Authors:** Alfred Chanda, Angela Gono-Bwalya, Lavina prashar

## Abstract

**Ethnopharmacological relevance:** Pregnant women in Chongwe, Zambia traditionally use the root of *Azanza garckeana* (F.Hffm.) Exell & Hillc (Malvalceae) to induce or accelerate labour. A previous study on the plant showed the crude root extracts to possesses uterotonic potential on isolated Wistar rat uterine smooth muscles. The methanol crude root extract was found to be the most potent.

**Aim:** The aim of the study was to isolated by bioactivity-guided isolation method the compound with the major uterotonic activity in the root of *Azanza garckeana*.

**Materials and methods:** This was a laboratory-based study designed using the uterus isolated from estrogenised adult non-gravid female virgin Wistar rats. The fresh plant materials (leaves and roots) were collected from Chongwe district. The methanol crude extract was obtained by continuous maceration and it was partitioned using hexane, chloroform, ethyl acetate and n-butanol in the increasing order of polarity. The most active fraction was fractionated using column chromatography. One-way ANOVA, Bonferroni post hoc tests were used to analyze data using STATA version 13. Bar charts and dose-response curves were done using Graphpad Prism version 5.00 for Windows (San Diego California USA).

**Results:** The final aqueous suspension demonstrated the highest uterotonic potency (EC_50_ = 2.49 x 10^−3^mg/ml; 95% CI 1.19 x 10^−3^ to 5.23 x 10^−3^ mg/ml, p=0.0001), while sub-fraction pool number 2 (subfractions number 41 to 61) demonstrated the highest uterotonic potency (EC_50_ at 2.05 x 10^−3^mg/ml; 95% CI 2.09 x 10^−3^ to 3.85 x 10^−3^ mg/ml, p=0.0001). Sub-fraction pool number 2 was suggested to contain the compound with the major uterotonic activity which was suggested to be related to the family of glycosides after High Performance Liquid Chromatography (HPLC) and Fourier Transform Infrared Spectroscopy (FTIR) analysis.

**Conclusions:** The study suggests the presence of a major uterotonic phytochemical constituent in the methanol root crude extract of *Azanza garckeana*, which was suggested to be related to the family of glycosides. The study has provided scientific evidence suggesting the root of *Azanza garckeana*, a plant used traditionally for inducing or accelerating labor possess uterotonic activity. Further pharmacological and toxicological studies need to be undertaken on the plant.

## 1.0 Introduction

Research involving traditional and herbal medicines, uses pharmacological and chemical methods to screen plants for biological activity and to isolate their active phytochemical compounds. Active phytochemical compounds are then used as models or scaffold compounds for drug design or synthesis (Koparde *et al*, 2019). Crude plant extracts have been suggested to induce uterine smooth muscle contractions. The screening of these plants for uterotonic properties have been done in different countries using animal models both *in vivo* and *in vitro* (Nworu *et al*, 2007; Amiera *et al*, 2014; Bello *et al*, 2010; Watcho *et al*, 2010; Lwiindi *et al*, 2015).

Uterotonic activity is the ability of an agent to induce contractions or to increase the tonicity of uterine smooth muscles. The uterus is a thick, pear-shaped, muscular organ approximately, 7cm long, and 4–5 cm wide at its widest point. The myometrium is the middle muscular layer that makes up the major proportion of the uterine body. Smooth muscle contraction by different agonists results in a rapid increase in intracellular calcium. Intracellular calcium binds to four binding sites of calmodulin causing a conformational change which allows the calmodulin-calcium complex to interact and activate the inactive myosin light chain kinase (MLCK). MLCK rapidly phosphorylates the 20 kDa myosin light chain (MLC). This leads to conformational changes in the myosin head that causes actin activation of myosin ATPase, resulting in force generation and contraction of the muscle fibres (Goodman *et al*, 2010). Myometrial contraction and relaxation is influenced by cell membrane receptors such as prostaglandin, oxytocin, beta 2, muscarinic and histamine receptors.

Different types of uterotonic herbal plants are used to induce or accelerate labour by pregnant women in sub-Sahara Africa; this is especially common in rural areas and low-income populations (Tripathi *et al*, 2012). The uterotonic activity of some of these plants have been screened using *in vivo and in vitro* animal models with positive results. Few studies have been done to isolate uterotonically active phytochemical compounds from these plants (Gruber &O’brien, 2010).

In Zambia, pregnant women use *Azanza garckeana* (F.Hffm.) Exell & Hillc (family Malvalceae) to induce or accelerate labour (Maluma *et al*, 2017). A study by Chanda *et al* (2020), to screen *Azanza garckeana* crude root extracts for uterotonic activity on isolated Wistar rat uterine smooth muscles found the methanol crude root extract to be the most potent followed by the hot aqueous and cold aqueous extracts, respectively. We, therefore, set out to do a bioactivity-guided isolation of the methanol crude root extract of *Azanza garckeana* on isolated nongravid estrogenised Wistar rat uterine smooth muscles. The findings of this study provide scientific evidence on the major uterotonic constituent in the crude root extract of *Azanza garckeana*.

## 2.0 Methodology

### 2.1 Study design

The Experimental was designed using nongravid estrogenised female virgin Wistar rats which were sacrificed by cervical dislocation on the day of the experiment. the uterus horn was isolated, cut into longitudinal strips and suspended in the 25ml organ bath (AD Instruments). The uterine smooth muscle strips were connected to the signal transducer which was connected to the power lab (AD Instruments) and computer installed with Lab tutor software. The baseline contractions and Oxytocin were used as negative and positive controls, respectively. The experiments were carried out in triplicates (n=3).

### 2.2 Data collection tools

The quantitative data were collected using the Lab-tutor software designed to collect data on the amplitude of contraction produced during the experiment. The data collection tables were used to collect data on the various concentrations applied to the organ bath and to record the corresponding amplitude of contraction produced.

### 2.3 Materials and methods

#### 2.3.1 Azanza garckeana plant collection and identification

The fresh leaves and roots of the plant were collected from Chongwe district of Zambia (15°16’17.3’ South 28°45’53.0’ East) with the aid of a known local herbalist. The plant was taken for identification at the University of Zambia and its specimen (accession number of 22209) was deposited in the Herbarium (UZL).

#### 2.3.2 Azanza garckeana Methanol crude root extract preparation

*Azanza garckeana* roots were washed clean, cut into small pieces with a Laboratory axe, shade dried for 14 days in a well-ventilated place and crushed with a blender (1.75Litres Satin Russell hubb Blender). The crushed root materials were packed in airtight Ziploc plastics and stored in a refrigerator at 4 °C until required. The roots weighing 3kg were extracted using methanol solvent by continuous maceration using a magnetic stirrer for 72hours hours, and then the extract was stippled and filtered with Whitman filter paper (No. 1). The Methanol crude extract was then dried in the vacuum at the temperature of 40ºC to obtain the powder whose yield was calculated. The extract was put in Ziploc containers and stored in the refrigerator at 4 °C until required.

### 2.4 Experimental Animals (Specimen)

Animals were housed in the animal house at the University of Zambia, School of Medicine. Female Wistar rats were separated from male rats after birth as soon as they could be identified as female. The selected rats were kept in plastic cages at room temperature and on a 12 h light/dark cycle with access to pellet food and water ad libitum. The adult female virgin Wistar rats weighing from 150 to 200g and aged from 5 to 6 months old were used as specimens for the experiment (Chanda *et al*, 2020).

### 2.5 Isolated Wistar rat uterine smooth muscle mounting

24hours before the experiment, the Wistar rats were pre-treated with 0.2mg/kg Diethylstilbestrol. On the day of the experiment, the rats were sacrificed by cervical dislocation, uterus horns were dissected out, cleaned of excess fat and connective tissues, and cut into longitudinal strips of about 2cm. The uterine smooth muscle strips were suspended in the 25ml organ bath (AD Instruments) using a cotton thread in the organ bath containing De Jalon’s physiological solution (9g/l of sodium chloride, 0.5g/l Sodium hydrogen carbonate, D. Glucose, 0.402g/l potassium chloride and 0.08g/l hydrated calcium chloride). The suspended uterine smooth muscle strips suspended in the organ bath were maintained at 37°C and aerated with a mixture of 95% Oxygen (O_2_) in 5% Carbon dioxide (CO_2_) using an aquarium air pump (Model No: 9905). The uterine smooth muscle strips were connected to the signal transducer which was connected to the power lab (AD Instruments) and computer installed with Lab tutor software. The tissue tension was adjusted using the transducers to the resting uterine smooth muscle contractions of 5mN, and then the force of contraction was zeroed (0mN) using the PowerLab. The suspended uterine smooth muscle strips connected to the transducers were allowed to equilibrate for at least 30 minutes to obtain the baseline uterus contractions before the samples were applied to it.

### 2.6 Uterotonic evaluation

#### 2.6.1 Standard drugs used for the experiment

The drugs that were used in this study are as follows;

1. **Diethylstilbestrol** (Kunj Pharma pvt. Ltd) was used in this study to promotes thickening of the adult female Wistar rat’s uterine endometrial layer so that the uterus horns can be easily isolated from the pelvic cavity.
2. **Oxytocin** (Mylan Health pty Ltd, Australia) was used in this study as a positive control for the screening of fractions and sub-fractions for uterotonic activity.

#### 2.6.2 Exposure assessment

Non-cumulative doubling concentrations of fractions and sub-fractions of *Azanza garckeana* and standard drugs were added one at a time to the De Jalon’s physiological solution in the organ bath where the uterine smooth muscle strips were suspended. Each sample was allowed to act for 10 minutes and the amplitude of contraction was measured. The experiment was done in triplicate (n=3) for each sample.

### 2.7 Bioactivity-guided isolation of the bioactive compound with the highest uterotonic activity in Azanza garckeana

The Bioactivity-guided isolation was done by separating *Azanza garckeana* phytochemical compounds according to the order of polarity using solvent partitioning method. The methanol crude root extract was reconstituted with water and it was partitioned with n-hexane, chloroform, ethyl acetate and n-butanol, respectively. After solvent partitioning, the most potent fractions were chromatographically fractionated to obtain 2 plant sub-fractions one of which contained the major uterotonic compound (Figure 1).

**Figure 1.**
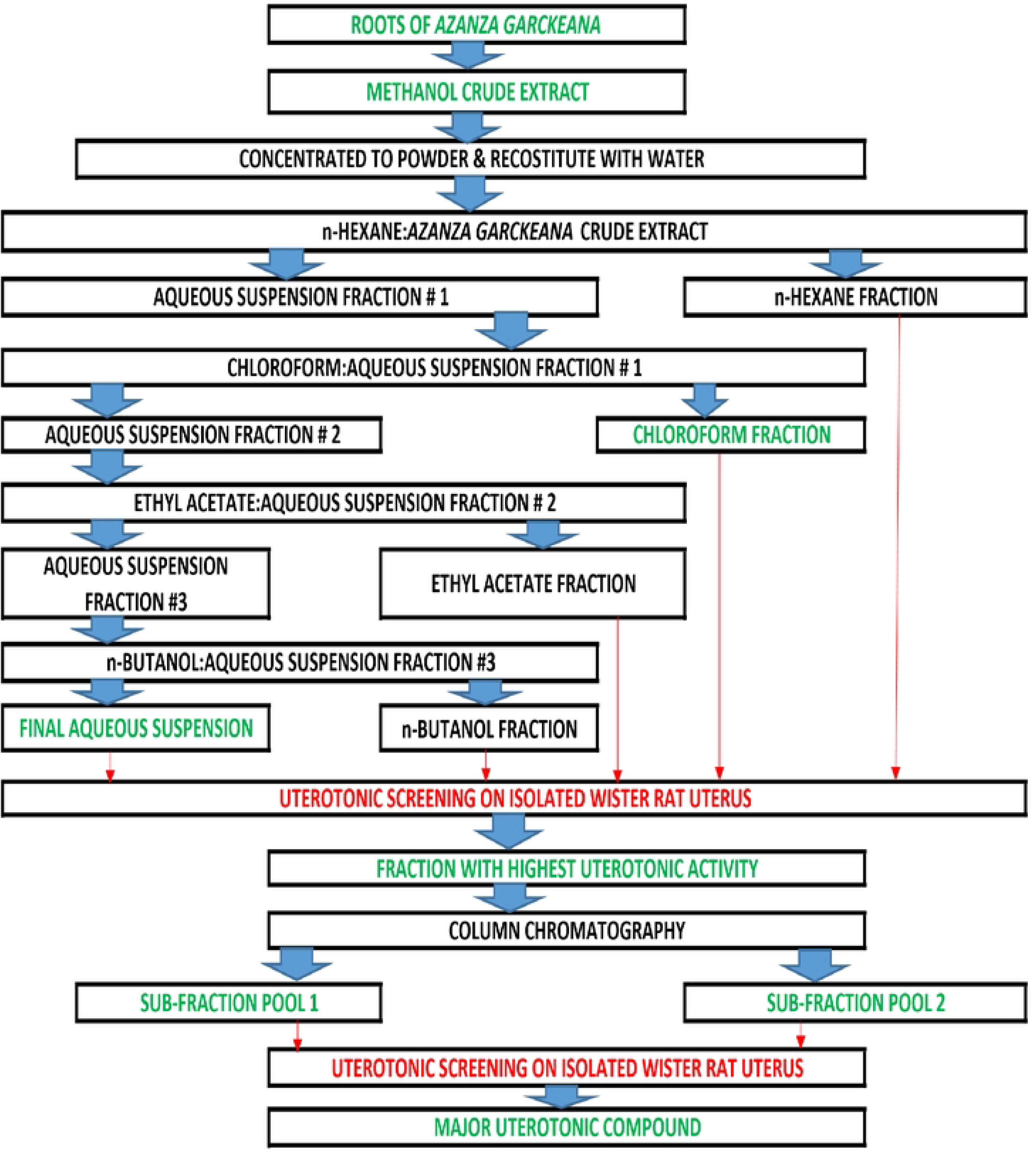
Stages of the Bioactivity-guided isolation process of Azanza garckeana methanol crude root extract on isolated Wistar rat uterine smooth muscles. The portioned solvents were agitated continuously for 2 minutes and were allo;ved to settle for 60 minutes to form 2 layers. The two layers of fractions were separated using a separating funnel. All fractions and sub-fractions were dried to powder before being screened for uterotonic activity.Thin Layer Chromatography (TLC) was done for each sub-fraction, and sub-fractions with similar TLC profiles (spots) were pooled. Sub-fraction pool one was obtained by pooling sub-fraction numbers 4 to 12 and 70 to 82. while Sub-fraction pool 2 resulted from the pooling of sub-fraction numbers41 to 61.

#### 2.7.1 Solvent partitioning fractionation

The powder of the methanol crude root extract was reconstituted with distilled water and was partitioned with organic solvents in the increasing order of polarity using a separating funnel. The crude extract with the highest activity was partitioned 3 times with 200ml of hexane, chloroform, ethyl acetate, and n-butanol (HIMedia Laboratories pvt. Ltd, India) in the increasing order of polarity to help with the fractionation of *Azanza garckeana* phytochemical compound as stated in Satyajit *et al* (2006) and Sudha & Srinivasan (2014).

#### 2.7.2 Preparation of the hexane fraction

To separate the nonpolar phytochemical compounds from the crude extract, it was reconstituted with 200ml of distilled water, partitioned with hexane (3 x 200ml), and was agitated continuously for 2 minutes and was allowed to settle for 60minutes to form 2 layers. All non-polar compounds, such as lipids and chlorophyll were in the hexane fraction (layer). This process is sometimes referred to as ‘‘defatting”. The hexane fraction was dried under vacuum to powder and was screened for uterotonic activity.

#### 2.7.3 Preparation of the chloroform fraction

The remaining aqueous suspension layer was partitioned 3 times with 200ml chloroform which was carried out by agitating the mixture continuously for 2 minutes. The mixture was allowed to settle for 60minutes to form 2 layers. Less polar phytochemical compounds were in the chloroform fraction, which was dried to powder and was then screened for uterotonic activity. The more polar compounds were in the remaining second aqueous suspension layer, which was carried forward for the next procedure.

#### 2.7.4 Preparation of the ethyl acetate fraction

The second aqueous suspension layer from the previous procedure was partitioned 3 times with 200ml ethyl acetate. This was done by agitating the mixture continuously for 2 minutes and allowed to settle for 60 minutes to form 2 layers. Less polar phytochemical compounds were in the ethyl acetate fraction, which was dried to powder and was then screened for uterotonic activity. The remaining third aqueous suspension layer was carried forward for the next procedure.

#### 2.7.5 Preparation of the n-butanol fraction

The third aqueous suspension layer from the previous procedure was partitioned 3 times with 200ml n-butanol. This was done by agitating the mixture continuously for 2 minutes after which it was allowed to settle for 60minutes to form 2 layers. More polar phytochemical compounds were in the n-butanol fraction, the fraction was dried to powder and was then screened for uterotonic activity. The remaining final aqueous suspension layer, which contained the most polar phytochemical compounds in *Azanza garckeana* was dried to powder and was then screened for uterotonic activity.

### 2.8 Silica gel filtration of Azanza garckeana fractions

The potent fractions of *Azanza garckeana* were further fractionated by silica gel column chromatography (silica gel: HIMedia Laboratories pvt. Ltd, India) to obtain sub-fractions. The Thin Layer Chromatography (TLC) was done for each sub-fraction, and the sub-fractions with similar TLC profiles (spots) were pooled.

### 2.9 Thin Layer Chromatography (TLC)

The Thin Layer Chromatography was conducted on a glass sheet coated with a thin layer of adsorbent material, which, in this case, was silica gel impregnated with a fluorescent material. Each component on the TLC appeared as spots which were separated vertically, and each had a retention factor (R_f_).

### 2.10 Analysis of the compound with the highest uterotonic activity

The isolated major uterotonic phytochemical compound was analyzed using High Performance Liquid Chromatography (Shimadzu, Japan). Ultraviolent (UV), wavelength 220nm was used as a detector for High Performance Liquid Chromatography (HPLC), the mobile phase used was Acetonitrile: Water (28:72) and the column used was C18 2.7µm, 3.0x100mm (Cortecs part number 1860007372). The HPLC oven temperature was at 20°C and the flow rate used was 0.03ml/min. The isolated compound was also analyzed using Fourier Transform Infrared Spectroscopy (Shimadzu, Japan) to identify the functional groups present on the compound. The wavenumbers (1/cm) obtained from the Fourier Transform Infrared Spectroscopy (FTIR) spectra were compared to the standard wavenumbers to help identify the suggested present functional groups on the isolated compound (Socrates, 2004; Ognyanov et al, 2018).

### 2.11 Display of data

In this study, data was displayed using tables, dose-response curves (concentration vs. amplitude of contraction) and figures. All data was presented as mean±standard error of the mean (SEM), 5% level of significant and 95% confidence interval was also displayed.

### 2.12 Data analysis

The primary variable was concentration in milligrams per milliliter (mg/ml) while the secondary variable was the amplitude of contractions in millinewtons (mN). To determine the differences in uterotonic activity within the various concentrations, one-way ANOVA was used. Bonferroni post hoc test was used to test at which concentration significant contractions were observed. STATA version 13 and Graphpad Prism version 5.00 for Windows (San Diego California USA) were used to analyze data. Origin software was used for FT-IR spectra. 5% level of statistical significant and 95% confidence interval were used.

### 2.13 Ethical considerations

The animals were treated humanely and were given access to standard nutrition, water and environment. The animals were sacrificed by cervical dislocation before the isolation of the uterus.

The study was conducted according to the guidelines for the design and statistical analysis of experiments using laboratory animal (Festing & Altaman, 2018; Council for International Organization of Medical Sciences and the International Council for Laboratory animal science, 2012). Approval was sort and obtained from the University of Zambia Biomedical Research Ethics Committee (UNZABREC) before the study was conducted (**REF. No. 006-12-18)**.

## 3.0 RESULTS AND DISCUSSION

### 3.1 Description of the Azanza garckeana methanol extracts

The crude root methanol extract of *Azanza garckeana* was dark brownish-black in colour with a percentage yield of 6.51% the finding which was corresponding to a similar study were the methanol crude root extract had the percentage of 6.26%, but was lower than the hot aqueous (22.26%) and cold aqueous (11.32%) extracts (Chanda *et al*, 2020). Alawode *et al* (2020) looked at the effect of extraction solvents on the phytochemical yield of the pulp and shaft of the plant and reported the methanol extract to have the highest percentage yield. The variations in the percentage yield for the 3 studies could be due to differences in geographical location and the plant parts used during the extraction method.

### 3.2 Solvent partitioning of the Methanol crude root extract of Azanza garckeana

The methanol crude root extract was selected for solvent partitioning because a previous similar study demonstrated it to be the most potency extract (EC_50_ = 1.28 x 10^−2^ mg/ml) as compared to the hot (EC_50_ =0.02792mg/ml) and cold (EC_50_ =0.4884mg/ml) aqueous extracts. The findings on the effect of *Azanza garckeana* crude root extracts on isolated Wistar rat uterine smooth muscles may suggest the Methanol crude root extract to contain the highest concentration of the phytochemical compound with the major uterotonic activity (Chanda *et al*, 2020).

### 3.3 Uterotonic activity of Azanza garckeana fractions

The amplitudes of contractions produced by the Hexane fraction (p= 0.2045), ethyl acetate fraction (p=0.2341) and n-Butanol fraction (p=0.5847) were not statistically significant. The amplitude of contraction produced by the chloroform fraction (p = 0.0001) and final aqueous fractions (p <0.0001) were statistically significant. The final aqueous fraction was not only more potent (EC_50_ = 2.56 x 10^−3^mg/ml; 95% CI 1.19 x 10^−3^ to 5.23 x 10^−3^ mg/ml, p<0.0001) than the chloroform fraction (EC_50_ = 1.09 x 10^−2^mg/ml; 95% CI 4.517 x 10^−3^ to 2.640 x 10^−2^ mg/ml, p=0.0001), but it was also more efficacious. The maximum amplitude of contraction produced by the fractions were 13mN for chloroform and 21mN for the final aqueous fraction, respectively (Figure 2).

**Figure 2:**
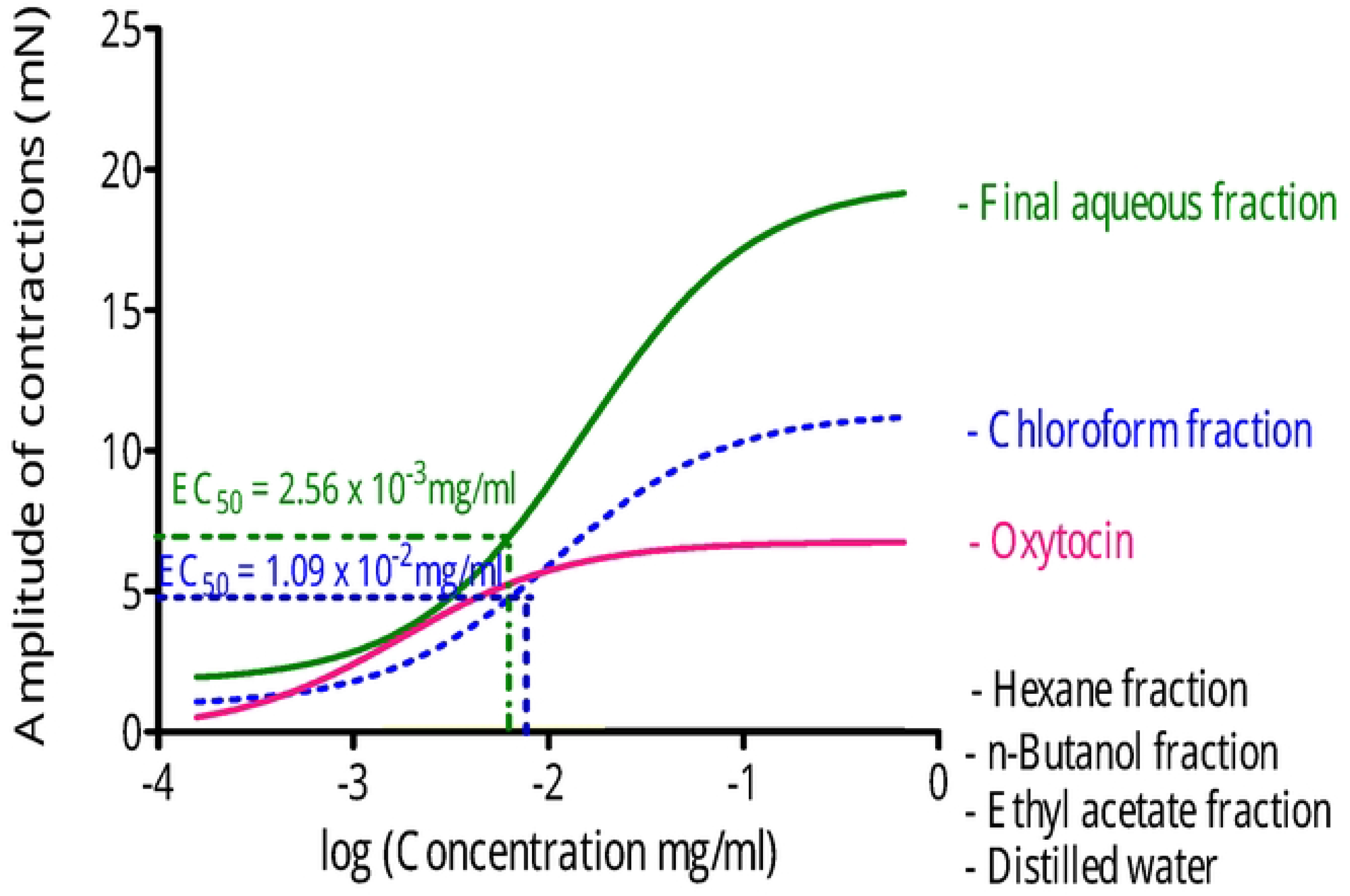
Concentration-dependent contraction of the uterine smooth muscle strips to 5 Azanza garckeana fractions. Points represent means_ SE M for the number of experiments (n=3). Distilled water, negative control; Oxytocin, positive control.

The activity of the Chloroform fraction started slow, but started to increase linearly after 2.56 x 10^−3^mg/ml and reached a plateau after 4.10 x 10^−2^ mg/ml. The activity of the final aqueous fraction increased linearly after 6.40 x 10^−4^ mg/ml and reached the plateau after 4.10 x 10^−2^ mg/ml. The t-test analysis of the potency for each of the *Azanza garckeana* fractions with that of Oxytocin also showed that the potency of the fractions was different from that of Oxytocin (Chloroform fraction EC_50_ 1.09 x 10^−2^ mg/ml, 95% CI 9.42 x 10^−4^ to 1.24 x 10^−2^ mg/ml, p<0.0001; Final aqueous fraction EC_50_ 2.56 x 10^−3^ mg/ml 95% CI 2.30 x 10^−3^ to 2.68 x10^−3^ mg/ml, p<0.0001).

The uterotonic verification of five fractions from the fractionation of the methanol crude extracts of *Azanza garckeana*, showed that only the chloroform soluble fraction and the final aqueous suspension fraction possessed significant uterotonic activity. The final aqueous suspension was more potent and efficacious than the chloroform fraction. This suggested that the compound with the highest uterotonic activity might have been in this fraction. When organic solvent partition fractionation is performed the final aqueous fraction contain glycosides with long sugar chains, hence the compound with the highest uterotonic activity was thought to be a glycoside which have also been previously isolated from *Azanza garckeana* (Satyajit *et al*, 2006). The final aqueous suspension demonstrated activity that was less potent than the standard uterotonic drug Oxytocin, but it demonstrated higher amplitudes of contraction at both the Emax and the EC_50_ concentrations. The higher amplitude of contraction produced by the final aqueous suspension fraction of *Azanza garckeana* suggested that the compound with the highest uterotonic activity may not have been lost during the solvent partitioning fractionation process.

The other fractions obtained from the methanol extract of *Azanza garckeana* were hexane, chloroform, ethyl acetate, and butanol fractions, respectively. The n-hexane fraction may have contained nonpolar compounds, such as lipids and chlorophylls. The process of removing fats and other non-polar compounds from a plant extract using n-hexane is sometimes referred to as defatting. After defatting, the remaining aqueous suspension may have been clear of non-polar compounds, which were inactive as far as the uterotonic activities were concerned. Less polar compounds may have been present in the chloroform soluble fraction. Polar compounds probably up to monoglycosides may have been present in the ethyl acetate soluble fraction. The n-butanol soluble fraction may have contained polar compounds which were mainly glycosides, and the final aqueous suspension fraction may have contained polar glycosides with polysaccharides (Visht & Chaturvedi, 2012; Satyajit *et al*, 2006).

### 3.4 Silica gel filtration of Azanza garckeana fractions

Thin Layer Chromatography (TLC) of the methanol extract of *Azanza garckeana* yielded three (3) spots which might suggest that the herbal plant might contain three (3) phytochemical compounds. The TLC results are in line with the findings of Nkafamiya *et al* (2015), who in their phytochemical screening of the root crude extract of *Azanza garckeana* using chemical methods found that the plant contained Alkaloids, Phenols and Saponins (Nkafamiya *et al*, 2015). It must be noted that although comparisons have been made to other studies, the TLC spots yield depend on factors such as the extract, mobile and stationary phases and the concentration of the sample applied to the TLC plate.

The amplitudes of contractions produced by the sub-fraction pool number 1 were significant (F =20.29, p <0.0001). The activities were significantly observed at the final bath concentration of 2.56 x 10^−3^mg/ml (p<0.0001). The sub-fraction had the EC_50_ at 1.47 x 10^−2^ mg/ml (95% CI 5.59 x 10^−3^ to 3.88 x 10^−2^ mg/ml) and the Emax at 4.10 x 10^−2^mg/ml. The amplitudes of contractions (mN) produced by the sub-fraction pool number 2 were significant (p <0.0001). The activities were significantly observed at the final bath concentration of 1.60 x 10^−4^mg/ml (p<0.0001). The pool had the EC_50_ at 2.05 x 10^−3^mg/ml (95% CI 2.09 x 10^−3^ to 3.85 x 10^−3^ mg/ml, p=0.0001) and the Emax at 4.10 x 10^−2^mg/ml. The pool of sub-fractions 41 to 61 which was named as sub-fraction pool number 2 was not only more potent than sub-fraction pool number 1 (pool of sub-fraction 4 to 12 and 70 to 82), but it was also more efficacious. Both pools of sub-fractions were less potent and efficacious as compared to the standard uterotonic drug Oxytocin (Figure 3).

**Figure 3.**
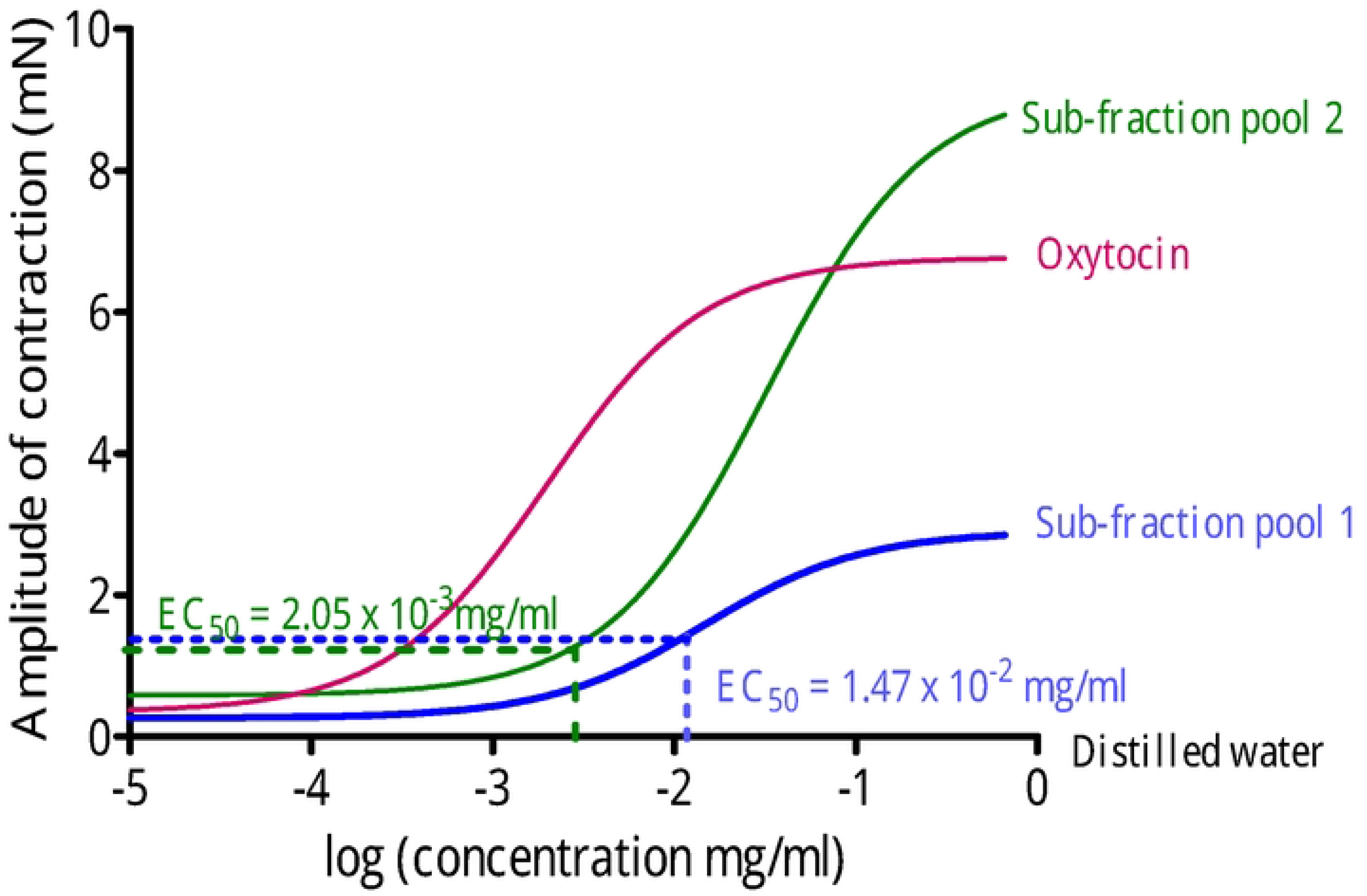
Concentration-dependent contraction of the uterine smooth muscle strips to sub-fraction pools 1 and 2. Points represent means_ SE M for the number of experiments (n=3). Distilled water, negative control; Oxytocin, positive control.

### 3.5 Analysis of the phytochemical compound with the highest uterotonic activity

#### 3.5.1 High Performance Liquid Chromatography of the isolated compound

The compound with the highest activity was isolated and was in the sub-fraction pool number 2 (sub-fractions number 41 to 61). The final aqueous suspension, which demonstrated the highest activity was suggested to be a polar glycoside with polysaccharides. The compound with the highest uterotonic activity might have been a glycoside, this was accessed by using a glycoside drug called digoxin as a UV-HPLC standard. The UV-HPLC chromatogram for both the isolated compound and the standard glycoside yielded similar peaks with the retention time of 16.729 minutes and 16.695 minutes, respectively.

#### 3.5.2 Fourier Transform Infrared Spectroscopy (FTIR) of the isolated compound

The functional groups of the isolated compound were identified by comparing the obtained wavenumbers to the standard FTIR wavenumber ranges. The FTIR spectra of the isolated compound had a similar FTIR spectrum with pectin as suggested by the FTIR feedback.

The Fourier Transform Infrared Spectroscopy (FTIR) spectra of the isolated compound with the highest uterotonic activity from *Azanza garckeana* showed the wavenumber at 3400 1/cm, 2900 1/cm, 1600 1/cm, 1400 1/cm, 1025 1/cm, 800 1/cm, 600 1/cm and 425 1/cm which might be due to the OH, -CHO, COO−, COO−/−OH, C-O-C), aromatic ring, C-H bending/Polygalacturonic acid and C-O-C torsion deformation in methyl polygalacturonate, respectively. The FTIR spectra for the major isolated uterotonic compound in *Azanza garckeana* was similar to that of pectin which is a structural hetero-polysaccharide hydrocolloid found largely in fruit and not in the roots of plants. Pectin are contained mainly in the primary cell walls of many plants and have previously been reported to possess various pharmacological activities (Ognyanov *et al*, 2018; Joel *et al*, 2018). Pectin has previously been isolated from the fruits of *Azanza garckeana* by Joel *et al* (2018) which had the FTIR spectra similar to the isolated uterotonic compound in this study. The FTIR spectra done on pectin by Joel *et al*. (2018) showed the broad band at 3415.19 cm-1 which could be due to SP hybridized C-H stretching vibration, 2935.40 cm-1 may probably be due to the SP3 hybridized C-H, 2380.32 cm-1 is likely to be C=C stretching vibration, the absorption band at 1639.02 cm-1 through 1737.56 cm-1 may probably be due to the C=O stretching vibration mode, 1533.85 cm-1 is likely due to the presence of N-H bending motion of the amine, 1380.87 cm-1 through 1460.29 cm-1 is due to SP3 hybridized CH2 of methylene bridge, 1059.13cm-1 through 1246.46 cm-1 is likely due to the C-O, and 769.51 cm-1 is due to the SP2 hybridized C-H bending vibrations. Pectin naturally occurs with various substituents such as ferulic acid a compound previously isolated from *Sida acuta* (Malvaleae). *Sida acuta* is a member of the Malvaleae family, a family from which the plant *Azanza garckeana* belong to. Abat *et al* (2017) reported that Ferulic acid is an active substituent which possesses other pharmacological activities such as anti-aging and Antidiabetic activities (Abat *et al*, 2017).

Carboxamine group is an essential substituent for the activity of uterotonic drugs such as oxytocin and Ergometrine, but ferulic acid does not contain the carboxamine substituent group in its structure and this can lead to the assumption that the isolated compound with the highest uterotonic activity maybe a pectin with the amidated ferulic acid substituent called N-Feruloyltyramine which has the uterotonically active Carboxamine group in its structure. N-Feruloyltyramine substituent is synthesized by the plant cell wall to resist or fight infection and may be attached to pectin in the cell wall of the plant.

The High Liquid chromatography (HPLC) analysis of the phytochemical constituent with the highest uterotonic activity suggests that it was structurally related to a glycoside called Digoxin which does not possess uterotonic activity. Guo *et al* (2008) suggested in their study that Pennogenin glycosides isolated from *Paris polyphylla* possess uterotonic activity. Pennogenin glycosides and Digoxin have similar structures which may suggest that the major uterotonic phytochemical compound in *Azanza garckeana* might be related to Pennogenin glycoside, but only full compound structure elucidation can ascertain this assumption (figure 4 and 5).

**Figure 4.**
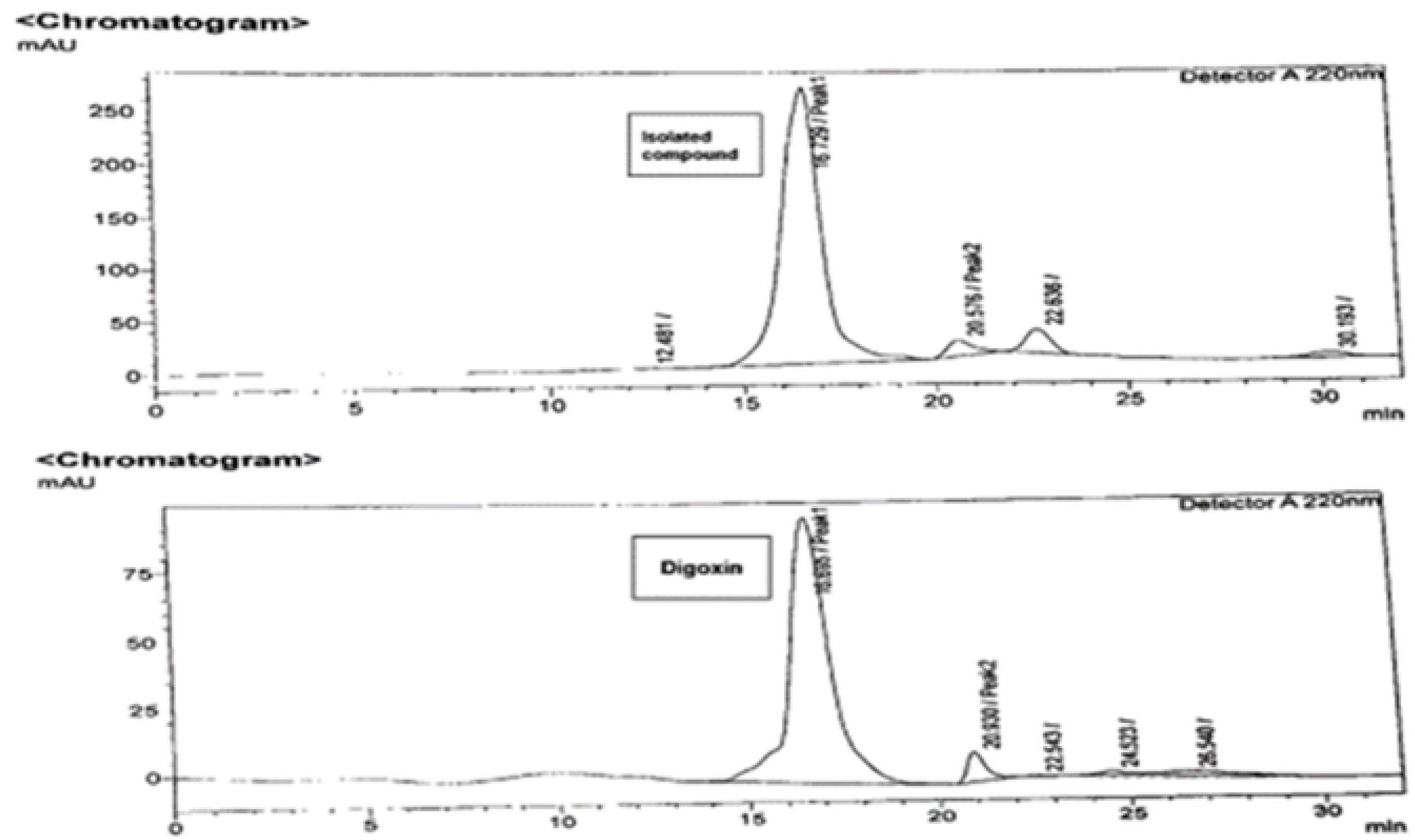
HPLC chromatogram for the isolated compound and Digoxin the standard glycoside.

**Figure 5.**
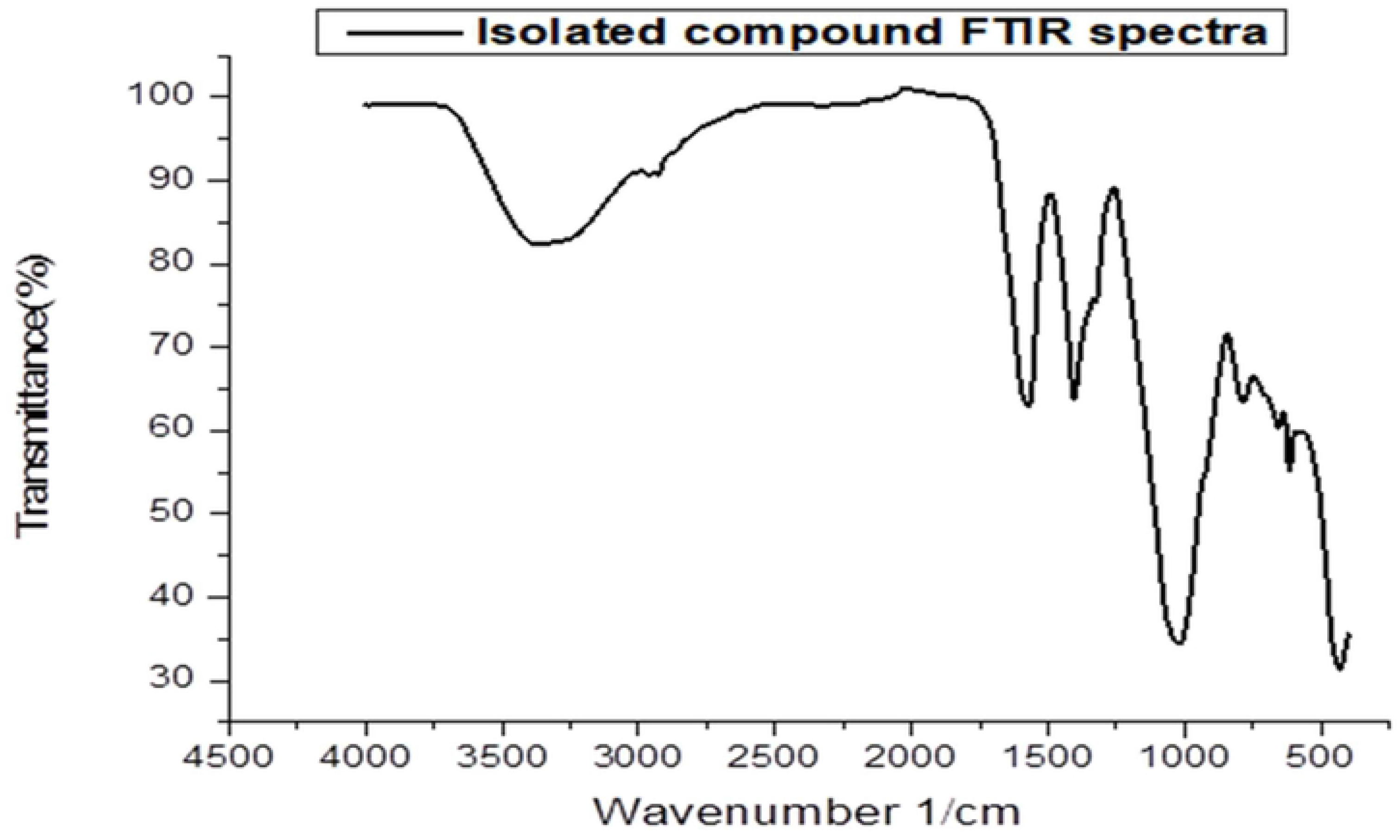
FTIR spectra for the isolated compound

## 4.0 CONCLUSION

The study showed that *Azanza garckeana* possesses uterotonic activity when evaluated using isolated Wistar rat uterine smooth muscles. The study suggests the presence of the major uterotonic phytochemical constituent in the methanol crude root extract of *Azanza garckeana*, which was suggested to be related to the family of glycosides. The study has provided scientific evidence suggesting that the root of *Azanza garckeana*, a plant used traditionally for inducing or accelerating labor possess uterotonic activity.

Further Pharmacological studies need to be done on the plant to confirm the phytochemical constituents responsible for the uterotonic effect of *Azanza garckeana*. The pharmacological studies should include the identification of all the phytochemical compounds responsible for the uterotonic activity of the plant and the structure-activity relationships of all these phytochemical compounds.

## 5.0 DATA AVAILABILITY

The data used to support the findings of this study are available from the corresponding author upon request.

## 6.0 ACKNOWLEDGEMENT

This study was partly supported by the Moses Sinkala research project Award.

## 8.0 AUTHORS’ CONTRIBUTIONS

AC developed proposal, collection data, analyzed data and prepared manuscript. AG provided scientific guidance and reviewed of Manuscript. LP provided technical support, scientific guidance, review of proposal and Manuscript.

## 9.0 ACKNOWLEDGEMENTS

The authors would like to thank the University of Zambia for laboratory access.

**Prof. MC Wilson and team** for their technical assistance and for awarding the research the Moses Sinkala research project ward, also to **Mr. Freddie Simwinga, Miss Mutinta, Mr. Golden Musamba and Mr. Newton Simfukwe** from the University of Zambia (UNZA), Ridgeway Campus for their valuable assistance during data collection. I would also like to thank **Mr. Siabamba** the chief Laboratory Technologist at the Chemistry Department, University of Zambia Great East Road Campus for his guidance and support during my data collection period.

## REFERECES

Abat, Jasmeet Kaur Kumar, Sanjay; Mohanty Aparajita. (2017). “Ethnomedicinal, Phytochemical and Ethnopharmacological Aspects of Four Medicinal Plants of Malvaceae Used in Indian Traditional Medicines: A Review” Medicines vol. 4, no. 4, pp 75.

Amiera Umi Romaizatul, Nihayah M, Wahida I. Farah, Raab N.F (2014). Photochemistry characteristic and uterotonic effect of aqueous extract of ficus deitoidea Leaves in Rats uterus. Pakistan Journal of Biological Sciences. Vol. 17, no.9, pp 1046–1051.

Alawode Ra, Ad Adesina, Ariyeloye Stephen Damola, Ia Mohammad et al (2020), Effect of drying methods and extractants on secondary metabolite compositions of Azanza garckeana pulp and shaft. Noble International Journal of Scientific Research. Vol. 02, no. 1, pp 01–07.

Chanda Alfred, Freddie Simwiga, Patrick Kaonga, Angela Gono-Bwalya, Lavina Prashar (2020). The Uterotonic Screening of the Root Extract of Azanza garckeana (Malvaceae) on Isolated Wistar Rat Uterine Smooth Muscles. Evidence-Based Complementary and Alternative Medicine. Vol. 2020, Article ID 8873180, pp.7. 10.1155/2020/8873180. Available online at https://www.hindawi.com/journals/ecam/2020/8873180

Bello SO, Bashar L, Muhammad BY, Onyeili P. (2010). Acute toxicity and uterotonic activity of aqueous extract of Lawsonia inermis (Lythraceae). Research Journal of Pharmaceutical, Biological and Chemical Sciences. Vol. 1, no. 3, pp.791.

Council for international organization of medical science (CIOMS) and the International council for Laboratory animal science (ICLAS). (2012). International guiding principles for biomedical research involving animals. Available online https://olaw.nih.gov/sites/default/files/Guiding_Principles_2012.pdf

Goodman L.S, Gilman A, Bruton L.L, Chabert B, Knollman B, (2010). Goodman and Gilman the Pharmacological basis of Therapeutics 12th Edition. The macGraw Hill companies,inc New york, USA. Chapter. 46, pp. 1339–1340.

Gruber W. Christian, O’Brian Margaret, (2010). Uterotonic plants and their bioactive constituents. Planta medica. Vol. 77, no. 3, pp. 207–220.

Guo L, Su J, Deng B W, Yu Z Y, Kang L P, Zhao Z H et al. (2008). Active pharmaceutical ingredients and mechanisms underlying phasic myometrial contractions stimulated with the saponin extract from Paris polyphylla Sm. var. yunnanensis used for abnormal uterine bleeding. Human Reproduction. Vol. 23, no. 4, pp. 964–971.

Joel J M, Barminas J T, Riki E Y, Yelwa J M, Edeh F (2018). Extraction and Characterization of Hydrocolloid Pectin from Goron Tula (Azanza garckeana) fruit. World Scientific News. Vol. 101, pp. 157–171.

Karas K, Sałkowska A, Sobalska-Kwapis M, Walczak-Drzewiecka A, Strapagiel D, Dastych J, Bachorz RA and Ratajewski M (2019) Digoxin, an Overlooked Agonist of RORγ /RORγ T. Frontiers in Pharmacology. Vol. 9, pp. 1460.

Koparde, A. A., R. Doijad and C. Magdum (2019). Natural Products in Drug Discovery. Available online at https://pdfs.semanticscholar.org/0249/45f5d18e3ac2aa5901b73a303732da9a4935.pdf?_ga=2.7538100.72471525.1610569535-1300532868.1586815982

Lwiindi L, Goma F, Mushabati F, Prashar L, Chongo K. (2015). The physiological response of uterine muscle to steganoteania araliacea in rat models. Journal of Medical science & Technology. Vol. 4, no. 1, pp. 40–45.

Maluma Sylvia, Kalungia A. Chichonyi, Hamachila Audrey, Hangoma Jimmy, Munkombwe Derrick. (2017). Prevalence of Traditional Herbal Medicine use and associated factors among pregnant women of Lusaka Province, Zambia. Journal of Pediatric Rehabilitation Medicine. Vol. 1, no. 1, pp. 5–11.

National Center for Biotechnology Information (2020). PubChem Database. Available online at https://pubchem.ncbi.nlm.nih.gov/compound/Pennogenin.

Nkafamiya L.L, Aldo B.P, Osemeahon S.A, Akinterinwa A. 201, (2015). Evaluation of nutritional, nontraditional, elemental content and amino acid profile of Azanza garckeana (Gordon Tula). British Journal of Applied Science & Technology. Vol. 12, pp. 1–10.

Nworu C.S, Akah P.A, Okoli C.O, Okoye T.C. (2017). Oxytocic activity of Leaf Extract of Spondia mombin. Pharmaceutical Biology. Vol. 45, no. 5, pp. 366–371.

Ognyanov M, Georgiev Y, Petkova N, Ivanov I, Vasileva I and Kratchanova M (2018), Isolation and Characterization of Pectic Polysaccharide Fraction from In Vitro Suspension Culture of Fumaria officinalis L. International Journal of Polymer Science. Vol. 2018. Available online at https://www.hindawi.com/journals/ijps/2018/5705036/

Satyajit D, Sarker Z, Latif A, Gray I (2006), Method in biotechnology TM _Natural Products Isolation, Second Edition. Chapter 10, pp. 269–273. Humana press, Totowa, new jersey. Available at https://www.humanapress.com

Socrates George (2004), Infrared and Raman Characteristic Group frequencies: Tables and Charts, 3rd Edition. John Wiley and sons Ltd, Chichester, New York, USA.

Sudha A and Srinivasan P (2014). Bioassay-guided isolation and antioxidant evaluation of flavonoid compound from Aerial parts of Lippia nodiflora L. available online at https://www.hindawi.com/journals/bmri/2014/549836/

Tripathi V, Stanton C and Anderson F W J (2012). Traditional preparations used as uterotonics in Sub-Saharan Africa and their pharmacologic effects. International Journal of Gynecology and Obstetrics. Vol. 120, pp. 16–22. Available xonline at 10.1016/j.ijgo.2012.06.020

Visht S and Chaturvedi S (2012). Isolation of Natural Products. Current Pharma Research. Vol. 2, no. 3, pp. 584–599.

Watcho Perrie, Ngagjui Esther, alango-efouet pepin, Nguelefack Telesphone Benoit, Kamanyi Albert (2010). Evaluation of in vitro uterotonic activities of fruit extract of ficus asperifolia in Rat. Evidence-Based Complementary and Alternative Medicine. Vol. 2011, pp. 1–7. Available online at https://www.hindawi.com/journals/ecam/2011/783413/

